# Reversible and gain modulation of neuronal responses and sensorimotor behavior by mid-infrared stimulation

**DOI:** 10.1101/2022.04.27.489693

**Authors:** Tong Xiao, Kaijie Wu, Yali Ding, Xiao Yang, Chao Chang, Yan Yang

## Abstract

Neuromodulation serves as a cornerstone for brain sciences and clinical applications. Mid-infrared stimulation (MIRS) has been recently reported to cause non-thermal modulation of brain functions. However little knowledge of mechanisms hampers its application. Here we bridge across ion channels, neuronal signals, and behavioral performances associated with sensorimotor transformation to provide evidence of how the alternation of neuronal activity by MIRS guides the change of behavioral performance in awake-behaving pigeons. We compared effects on visually-guided eye movements by applying MIRS and electrical stimulation (ES) in the pretectal nucleus lentiformis mesencephali (nLM). Distinct from ES, we found a specific gain modulation of MIRS to alter behavior in a manner of the strength of visual inputs. Our simultaneous extracellular recordings showed that MIRS can excite and inhibit the neuronal activity in the same pretectal neuron based on its ongoing sensory responsiveness levels in awake-behaving animals. We further applied computational simulations and found that MIRS can modulate the carbonyl group (-C=O) enriched on the potassium channel to resonate, and could affect action potential generation, alter neuronal responses to sensory inputs and then guide behavior. Our findings suggest that MIRS could be a promising approach for brain researches and neurological diseases, with gene free manipulation.

## Introduction

Neuromodulation has long been employed to treat patients with brain disorders and answer scientific questions on brain functions. It is consistently growing and evolving with innovations in technology. Deep brain stimulation, and transcranial electromagnetic stimulation are thought to effectively activate neuronal excitability and connectivity, and are applied commonly in clinical treatments. Optogenetic stimulation shows power to selectively excite and inhibit specific groups of neurons, but the genetic manipulation limited its clinical applications (Deisseroth, 2015). It would be of great significance to clinical setting and neuroscience research if a neuromodulation achieves reliable neural excitation and inhibition, with gene free manipulation.

Optical infrared neuronal stimulation is emerging as a potential neuromodulation because of the ability to deliver focused energy through tissue even without direct contact. Initial studies showed that near-infrared wavelength stimulation (NIRS) could excite neuronal responses in *vitro* (Izzo, et al., 2008; Albert, et al., 2012; Shapiro, et al., 2012; Entwisle, et al., 2016) and in *vivo* (Wells, et al., 2005; Wells, et al., 2007a; 2007b; Richter, et al., 2008; Xia, et al., 2014). Interestingly, there are also limited cases observed that NIRS could inhibit neuronal firing (Cayce, et al., 2011; Duke, et al., 2012; Duke, et al., 2013; Horváth Á, et al., 2020). Recently, there are rising evidence shed light on that mid-infrared stimulation (MIRS) with a specific wavelength could lead to dramatic changes: 1) *neuronal firing*. MIRS can exert non-thermal effects on ion channels, and lead to gain modulation of action potentials based on current injections in *vitro* brain slices (Liu, et al., 2021). MIRS can enhance neuronal spontaneous activities (Zhang, et al., 2021) and sensory responses (Tan, et al., 2021) in anesthetized animals. 2) *behavioral performance*. MIRS can effectively modulate behavior by accelerating associative learning in mice (Zhang, et al., 2021), and regulating startle responses in larval zebrafish (Liu, et al., 2021). Although our understanding of how MIRS neuromodulation impacts the brain is evolving, it is still left vacant to demonstrate how MIRS alters *neuronal firing* in brain network, and then how the alternation guides *behavioral performance* for lack of simultaneous recording in awake-behaving animals.

In this research, we aim to bridge across cellular and behavioral phenomena to provide evidence of how alternations of neuronal activity by MIRS regulate behavioral performances in awake-behaving pigeons. When birds fly, surrounding environment generates a large field of visual motion across the entire retina, known as “optic flow” (Gibson, 1951). Their eyes reflex produce optokinetic nystagmus (OKN) to maintain stabilization of image on the retina. OKN combines a close tracking of a moving field by pursuit eye movement in slow phases, and a rapid resetting back by saccadic eye movement in fast phases, producing a characteristic irregular sawtooth waveform. In birds, the pretectal nucleus lentiformis mesencephali (nLM, homologous to the nucleus of the optic tract) (Fite, 1985; McKenna and Wallman, 1985) is an essential encoder to process horizontal visual information (Winterson and Brauth, 1985; Wylie and Frost, 1996; Fu, et al., 1998; Wylie and Crowder, 2000; Cao, et al., 2004; Wylie, et al., 2018) and generate OKN eye movements (Fite, et al., 1979; Burns and Wallman, 1981; Gioanni, et al., 1983; Cao, et al., 2006; Yang, et al., 2008a; 2008b; Wylie, 2013; Ibbotson, 2017; Gutierrez-Ibanez, et al., 2018). Majority of nLM neurons become excited by visual motion in the temporal-to-nasal direction and prefer slow velocity, whereas other neurons are predominantly sensitive to the nasal-to-temporal, or vertical motion. It transfers visual information from retina directly to multiple brain areas for perception and motor control during self-motion, oculomotor and others (Cao, et al., 2006; Yang, et al., 2008b; Wylie, 2013; Ibbotson, 2017; Wylie, et al., 2018).

In this study, we applied MIRS in the pretectal nLM and revealed neuromodulation effects in awaking-behaving pigeons by evidence for several phenomena. First, we found a reversible and gain regulation of pursuit velocity of OKN eye movements depending on the strength of visual inputs. Second, we simultaneously recorded neuronal activities in the pretectal nLM, and found that MIRS could facilitate and suppress firing activity based on levels of neuronal responses. Third, we have applied computational simulations and found MIRS could preferentially enhance potassium permeability through K^+^ channels to alter action potential generation, which would modulate neuronal signals in brain network and guide sensorimotor responses.

## Results

### MIRS exerts gain modulation of pursuit depended on the strength of visual inputs

We introduced a large-field grating motion to pigeons. Vision-evoked OKN eye movements were recorded before, during and after ∼120sec MIRS application in the left pretectal nucleus (Fig. 1A-C). When pigeons viewed a grating motion of 8 deg/s in the temporo-to-nasal (T-N) direction (Fig. 1D top plots), they closely pursued moving gratings (Fig. 1E; velocity: 4.88±0.29 deg/s; duration: 1.89±0.26 s; amplitude: 8.44±1.06 deg; mean ± SEM, n=10) along the T-N direction, and then quickly saccade back to reset eye position. Once MIRS was turned on, animals significantly fasted their pursuit performances (velocity: 7.16±0.60 deg/s; Wilcoxon signed-rank test, p<0.01), but kept similar pursuit durations and distances (duration: 1.71±0.26 s; amplitude: 8.66±0.95 deg, p>0.05). Conversely, when pigeons were introduced a grating motion of 8 deg/s in the nasal-to-temporal (N-T) direction (Fig. 1D, bottom plots), animals tracked grating motion with far less effective pursuit eye movements (Fig. 1E; velocity: -0.89±0.09 deg/s; duration: 2.95±0.47 s; amplitude: -1.80±0.16 deg). When MIRS was turned on, animals again significantly fasted their eye movements to pursue in the N-T direction (velocity: -1.44±0.09 deg/s; Wilcoxon signed-rank test, p<0.01), without significant changes in pursuit durations and distances (duration: 2.75±0.39s; amplitude: -2.22±0.22 deg, p>0.05). There was asymmetry between the N–T and T–N OKN, which has been widely observed in lateral-eyed vertebrates (rabbits: Collewijn, 1969; pigeons: Zolotilina, et al., 1995 rats: Harvey, et al., 1997; mice: Kodama and du Lac, 2016). Although there were massive asymmetric sensorimotor responses, MIRS significantly facilitated pursuit velocities in slow phases evoked by 8 deg/s grating motion along both T-N and N-T directions, in the individual and group animals.

**Figure 1.**
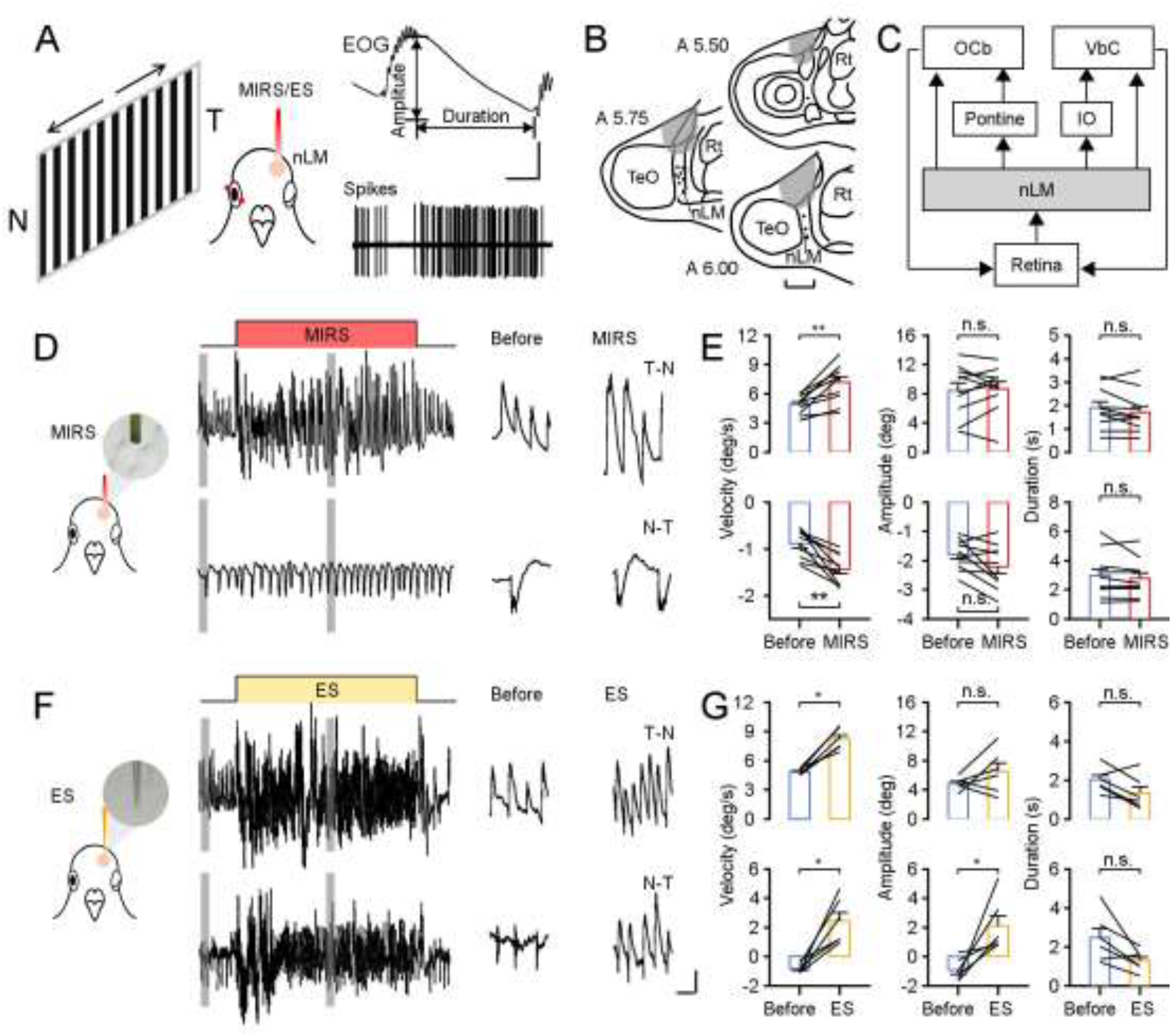
Modulation of sensorimotor behavior by MIRS in pigeons. **A:** Schematic drawing of neuronal activities and behavioral performances recording system, with visual stimuli of grating motions (T-N and N-T direction), from top to bottom: EOG traces of OKN, and action potentials of an example nLM neuron. **B:** Marked recording sites (dots) and MIRS/ES sites (gray shading) in the pretectal nLM cross brain sections under study (A 5.5-6.0). **C:** Optic flow pathways from nLM to the cerebellum for generation of OKN in birds. Abbreviations are: OCb, ocularmotor cerebellum; VbC, vestibular cerebellum; IO, inferior olive; nLM, the pretectal nucleus lentiformis mesencephali. **D, F:** MIRS (**D**) and ES (**F**) modulated OKN eye movements based on grating motion directions in example animals (top to bottom traces: T-N and N-T direction motion). **E, G:** Comparison of pursuit eye movement parameters to T-N and N-T grating motion (top and bottom plots) before and during MIRS (**E**, n=10) and ES (**G**, n=6). * P<0.05, ** P<0.01, Wilcoxon signed-rank test. Error bars represent SEM. Black lines represent data from individual animals. Scale bars in A: 0.3s, 12deg; B: 1mm; D and F: top-left 12s, 4deg; top-middle and top-right, 2s, 4deg; bottom-left 12s, 1deg; bottom-middle and bottom-right, 2s, 1deg.

Next, we conducted ES experiments with a classic frequency used in deep brain stimulation as a comparative research. Similarly, a grating motion of 8 deg/s was introduced to pigeons, and a ∼120sec ES was applied in the similar way as MIRS (Fig. 1F). Distinct from MIRS, ES effectively deflected eye movements to pursue toward the T-N direction, independent of visual motion directions of the T-N or N-T (Fig. 1F and G, T-N OKN in top plots, before: 4.84±0.12 deg/s; ES: 8.19±0.42 deg/s; N-T OKN in bottom plots, before: -0.80±0.14 deg/s; ES: 2.43±0.59 deg/s; mean ± SEM; Wilcoxon signed-rank test, n=6, p<0.05).

To verify differences in the modulation of MIRS and ES, we compared effects of MIRS and ES on pursuit movements of OKN. Pigeons sensitively produced OKN to large-filed grating motions of different directions and velocities, or spontaneously made saccades when they freely viewed stationary gratings (Fig. 2). When the grating moved at a faster velocity of 8 deg/s, MIRS significantly facilitated the pursuit eye movements along the grating motion direction. When the grating moved at a lower velocity of 2 deg/s or kept still (0 deg/s), changes of pursuit velocity by the same MIRS were insignificant (Fig 2A, T-N OKN, before: 1.97±0.05 deg/s; MIRS: 2.15±0.22 deg/s; N-T OKN, before: -0.40±0.11 deg/s; MIRS: -0.47±0.11 deg/s; spontaneous saccades, before: -0.09±0.07 deg/s; MIRS: 0.03±0.12 deg/s; Wilcoxon signed-rank test, p>>0.05). The effects of ES on OKN were consistent across different visual conditions: ES deflected pursuit eye movements towards the T-N direction, regardless of the visual motion information (directions or velocities of grating motion) (Fig. 2C, Wilcoxon signed-rank test, p<0.05). We further compared pursuit data that had been normalized to the top 10 trials with highest pursuit velocities before MIRS/ES under each visual condition. This could eliminate individual animal bias from population analysis. Data showed that gain modulation intensity of MIRS on pursuit depended on the strength of visual inputs (Fig. 2B and D): when the visual motion was “faster”, MIRS facilitated the pursuit eye movements of OKN significantly; however, when the visual motion was “slower”, MIRS failed to regulate oculomotor behavior.

**Figure 2.**
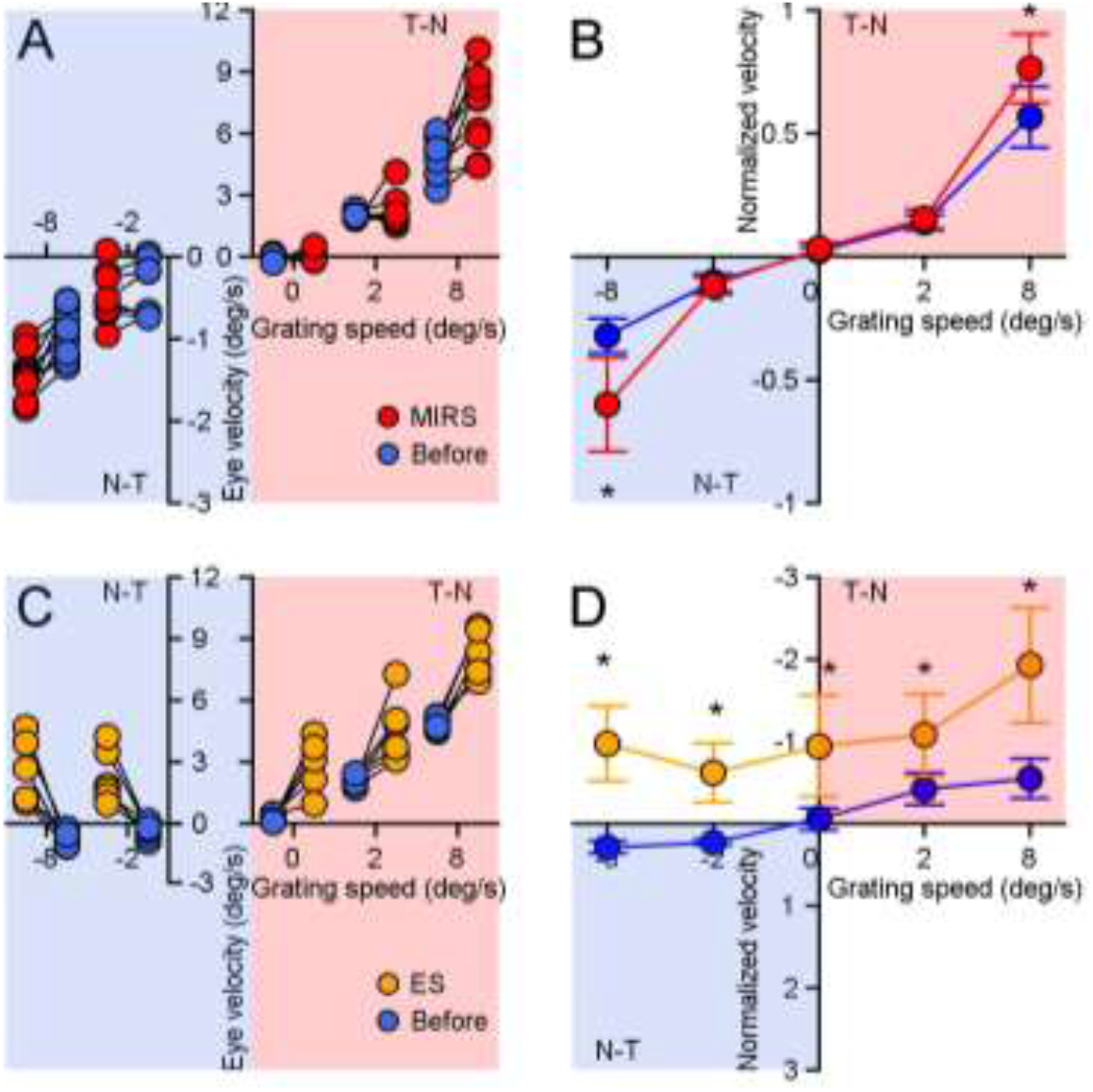
Comparison of modulation effects by MIRS and ES on OKN eye movements under different directions and velocities of grating motions. **A, C:** MIRS(**A**) and ES(**C**) modulated pursuit eye velocities based on grating motions. Black lines show data from individual animals (blue symbols: before stimulation; red symbols: during MIRS; yellow symbols: during ES). **B, D:** Pursuit eye velocity data had been normalized to average values from the top 10 trials with highest peak velocities before MIRS(**B**) and ES(**D**) under each visual input. Wilcoxon signed-rank test, * P<0.05. Error bars represent 1 SEM.

### MIRS excites and inhibits neuronal responses in the same pretectal neuron

To examine effects of MIRS on neuronal responses, we performed extracellular recording and tested 31 nLM neurons in awake-behaving pigeons before, during and after MIRS irradiation. 16 of them increased firing rates to a grating motion in the T-N direction (“preferred direction”), while decreased in the N-T direction (“null direction”). The other 15 neurons showed an oppositive direction selectivity. All recorded nLM cells have been modulated by corollary discharge signals during saccadic eye movements (Yang, et al., 2008a): their firing rates were inhibited during saccades. To investigate how nLM neurons coding sensory information to guide pursuit eye movements, we aligned the spiking activity to the onset of pursuit eye movements of OKN. The following data analysis focused on an interval from 0-600 ms after the onset of pursuit. Visual responses of 28 neurons were significantly modified during MIRS (Wilcoxon signed-rank test, p<0.01). Among them, only 3 neurons were suppressed firing rates under all conditions tested. We focused on the left 25 pretectal neurons for further data analysis.

An example nLM neuron with a preferred direction of N-T was shown in Figure 3. During MIRS, neuronal activities were modified by positive and negative effects linked with excitation and inhibition of visual responses. When the pigeon viewed a grating motion of 8 deg/s in the neuronal preferred direction (N-T), visual responses were excited to 31.09±6.80 spikes/s (mean ± SD). Once MIRS was turned on, neuronal excitation was further facilitated to 37.95±11.32 spikes/s (Student’s t-test, p<0.001, Fig. 3C and E) and fasted pursuit velocity toward N-T direction (Fig. 2A). When the pigeon viewed a grating motion in the null direction (T-N), neuronal responses were inhibited to 26.25±6.73 spikes/s. Interestingly, the same MIRS showed an oppositive effect and further significantly enhanced neuronal inhibition to 18.40±6.87 spikes/s (Student’s t-test: p<0.001, Fig. 3D and F), and speeded up pursuit eye movements to the T-N direction (Fig. 2A). Our data showed positive and negative modulations of neuronal responsiveness occurred in the same recorded neuron, depending on the level of visual responses.

**Figure 3.**
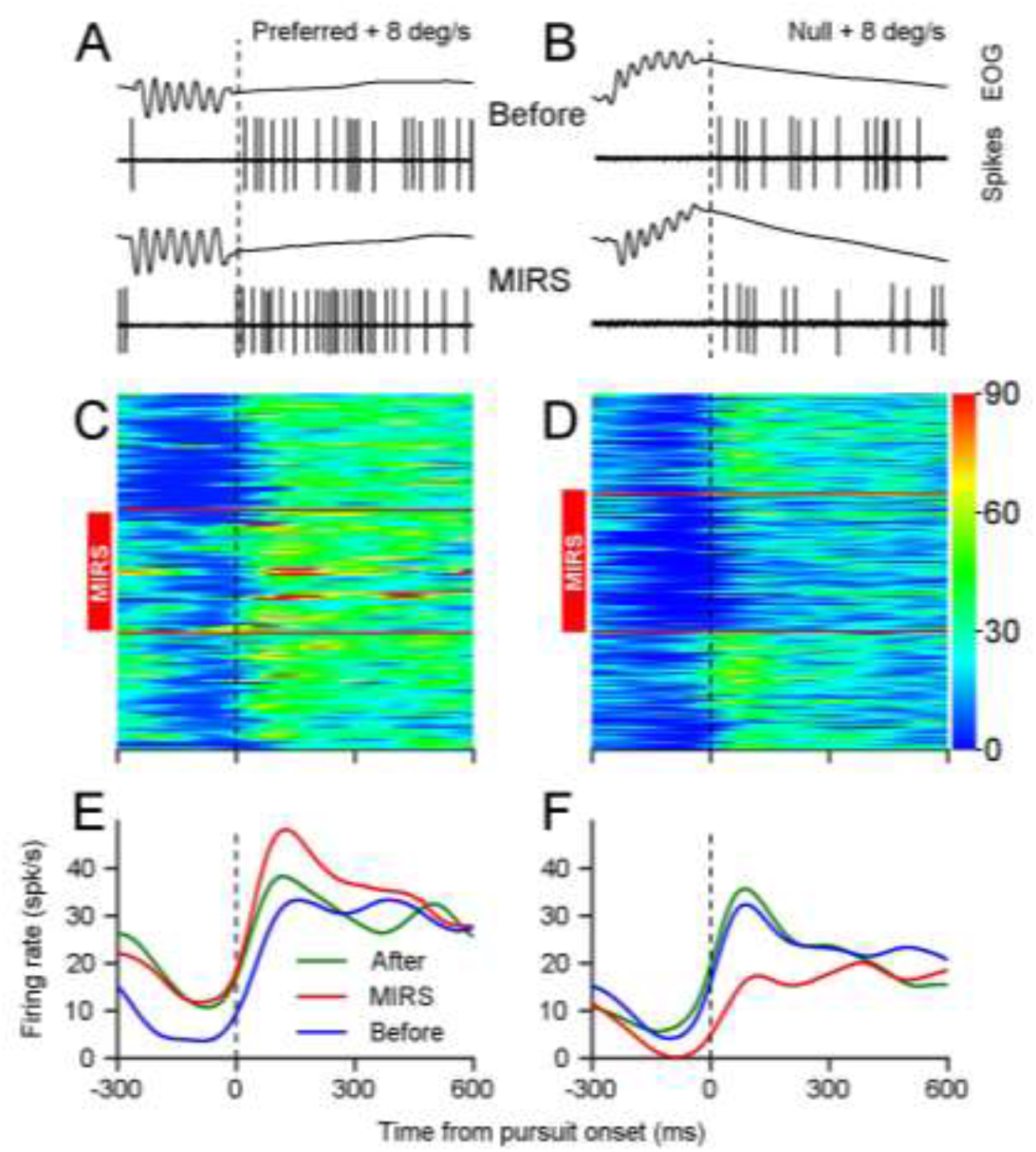
MIRS exerts positive and negative modulation of visual responses during pursuit in slow phases of OKN in an example nLM neuron. **A, B:** Representative eye movements and spiking responses to grating motion at 8 deg/s in the preferred (**A**) and null direction (**B**) of an example neuron. From top to bottom: EOG traces of OKN, and action potentials of the neuron before and during MIRS. **C, D:** Visual responses of the cell before, during and after MIRS are color-coded with a scale (spikes/s) on the right. MIRS facilitated neuronal excitation during pursuit to grating motion in preferred direction (**C**). The same MIRS enhanced neuronal inhibition during pursuit to grating motion in null direction in the same neuron (**D**). Each horizontal colored line shows neuronal firing rates during pursuit in one OKN. Red horizontal lines show the period of MIRS application. **E, F:** Mean firing rates of the example cell, as a function of time from the onset of pursuit. Blue, red and green traces show data obtained before, during and after MIRS, respectively.

Next, we compared neuronal population responses before, during and after MIRS while they were evoked by grating motions of 8 deg/s in preferred or null directions. All tested neurons are excited to their preferred direction (mean firing rates: 55.99 ± 24.02 spikes/s, n=21) or inhibited to their null direction (mean firing rates: 28.93 ± 24.19 spikes/s, n=14) before stimulations. We defined the ratio of firing rates by normalization to visual responses before MIRS for each cell. Our data suggested that MIRS could significantly facilitate neuronal excitation in the preferred direction and enhance neuronal inhibition in the null direction. These bidirectional effects of MIRS occurred reversibly (Fig. 4A-C, Wilcoxon signed-rank test, p<0.01).

**Figure 4.**
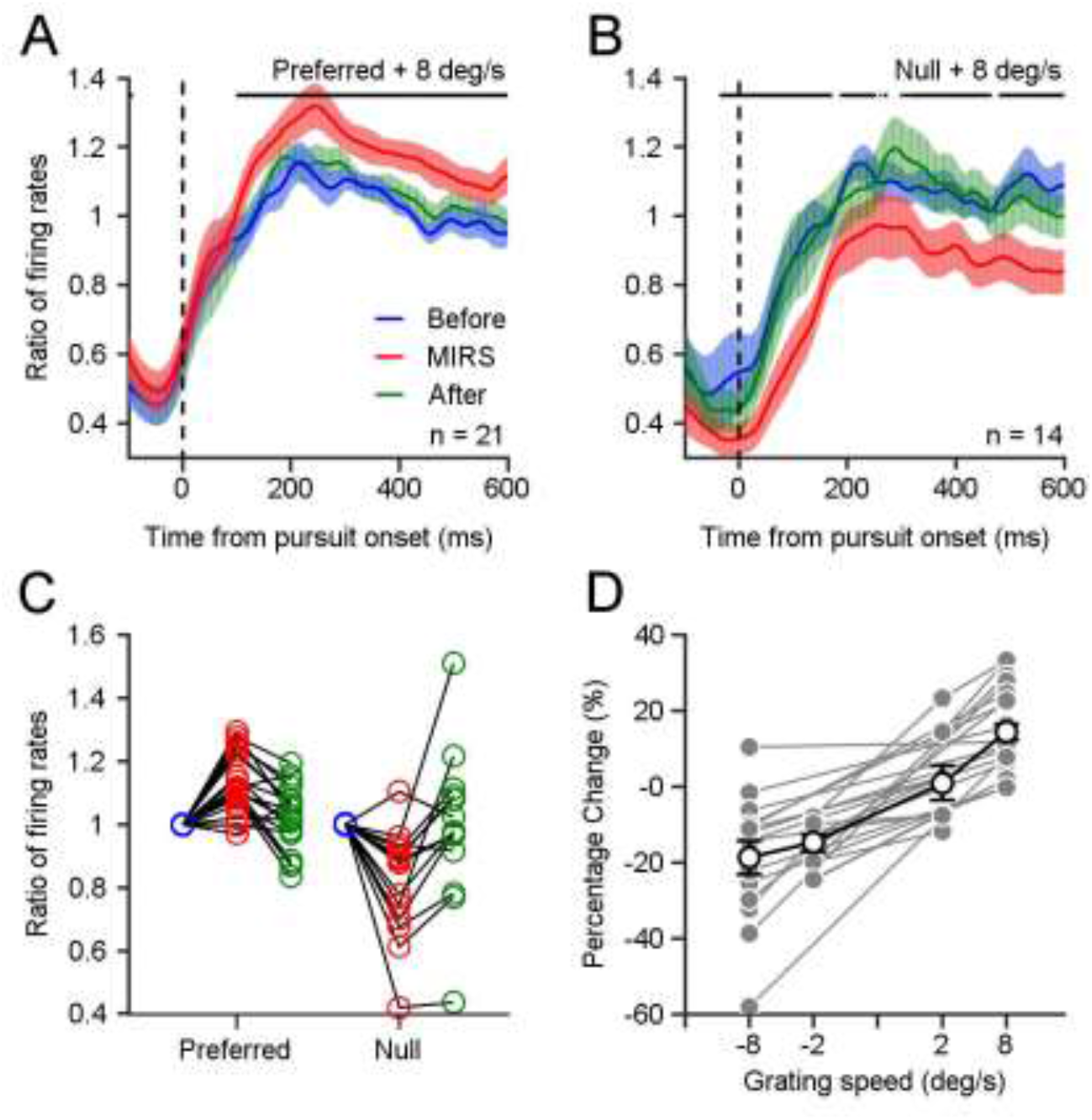
MIRS alters sensory coding in pretectal neurons associated with the level of visual responses. **A, B:** Comparison of neuronal population responses before, during and after MIRS while neurons were evoked by grating motions of 8 deg/s in the preferred (**A**) and null (**B**) directions. Data during and after MIRS were normalized to neuronal firing rates before stimulation. Black stars on the top showed that MIRS significantly increased(**A**) and decreased(**B**) neuronal responses (Wilcoxon signed-rank test, * P<0.05). **C:** Ratio of neuronal responses in A and B showing the effect of MIRS in preferred and null directions. Black lines show data from individual neurons. Blue, red and green symbols show data from before, during and after MIRS. **D:** The percentage change of visual responses across populations was correlated with the strength of visual inputs. Gray symbols and lines show data from individual neurons. Open symbols with black line show the average across populations. Error bars represent 1 SEM.

Further, we statically analyzed neuronal activities across population under different visual inputs. Pretectal nLM neurons’ firing rates can be evoked to different responsiveness by grating motions at 2 or 8 deg/s in preferred and null directions: when the grating motion speeded up from 2 deg/s to 8 deg/s in preferred or null directions, visual responses increased from ∼40 to ∼55 spikes/s or decreased from ∼25 to ∼15 spikes/s (Our unpublished data, also reported by (Cao, et al., 2004)). Visual inputs evoked neuronal initial discharges at different levels of high- or low-frequency. The percentage change of visual responses was correlated with the strength of visual inputs, even in individual neurons (Fig. 4D, each gray line presents the same tested neuron). When the grating moved at a higher velocity of 8 deg/s in the preferred direction, the percentage change of visual responses by MIRS (14.31% ± 2.10%, n=21) was larger than ones when the grating moved at a lower speed of 2 deg/s (0.84% ± 4.62%, n=7) and in null direction (−14.86% ± 2.32%, n=6 for 2 deg/s, Wilcoxon rank-sum test, p<0.01; -18.81% ± 4.43%, n=14 for 8 deg/s; Wilcoxon rank-sum test, p<<0.01). These results supported that MIRS exerts gain modulation of neuronal signals, in a manner that is itself sensory responses dependent.

### MIRS preferentially regulates permeation of K^+^ channels

To explore the potential molecular mechanism underlying the gain modulation of neuronal responses by MIRS, we constructed models of voltage-gated K^+^ and Na^+^ channels. Recently, models of biomimetic ion channels have been widely used to investigate the translocation events of K^+^ and Na^+^ ions (Long, et al., 2005; Zhang, et al., 2012). In our model (Supplementary Fig. S1), we constructed K^+^ channels containing the whole protein placed at the middle of phospholipid bilayer (DPPC molecules) to separate water and ions on each side, in which the model could provide a more authentic and informative simulation of the function of MIRS on ion channels. We started to define the absorption spectrum of ion channels to MIRS based on classical molecular dynamics (MD) method. The MD simulation showed a remarkable absorption finger of K^+^ channels located between 30 to 40 THz, while just out of the strong absorption spectral ranges by water molecules (Heyden, et al., 2010) and Na^+^ channels. The specific frequency of 34.88 THz applied in the study was closed to the maximum absorption spectrum, at least within half-height width of the finger (Fig. 5A).

**Figure 5.**
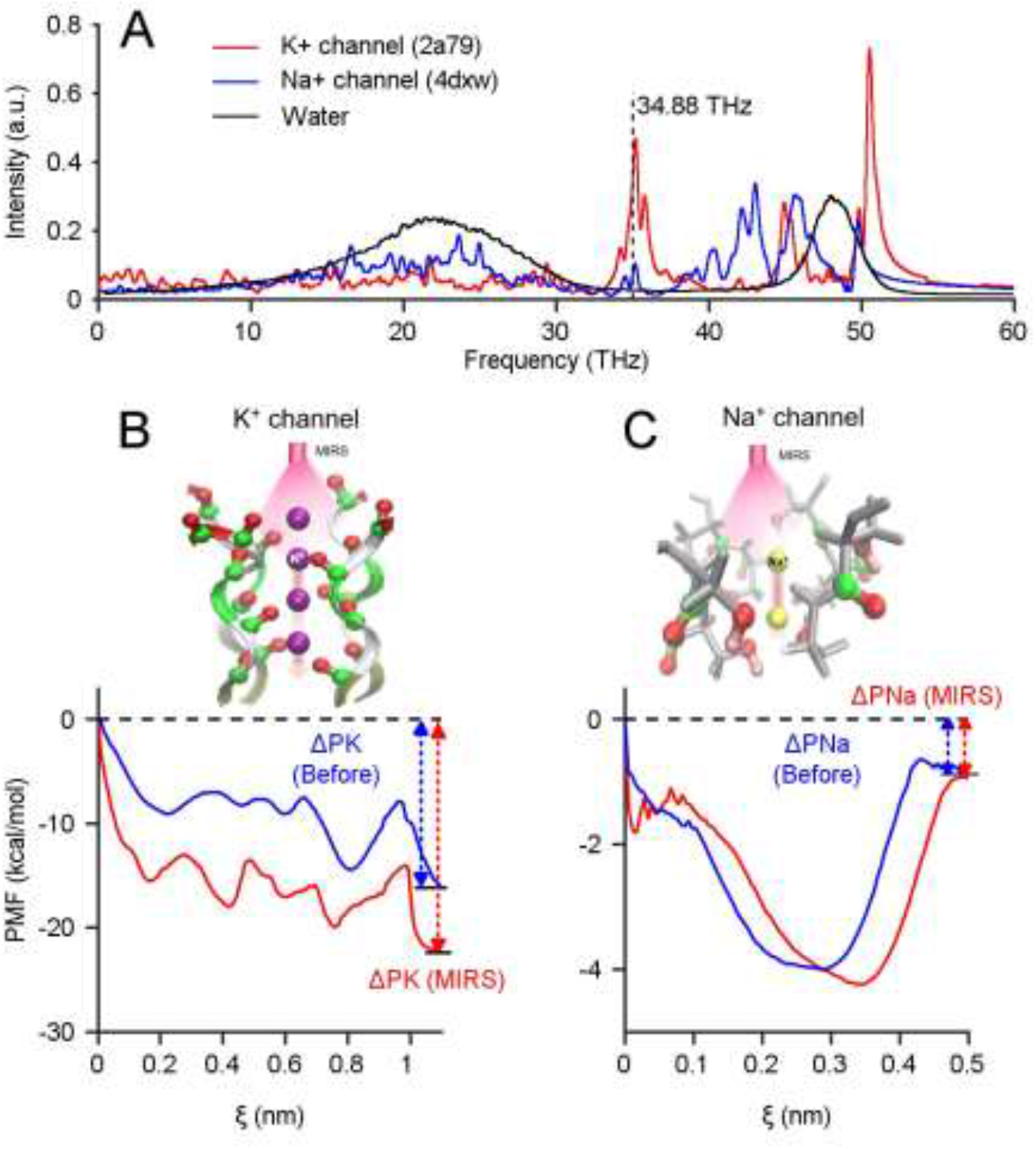
Computational simulations reveal preferential regulation of K^+^ channels by MIRS. **A:** Absorption spectra of K^+^ and Na^+^ channels were calculated by using molecular dynamics simulation, respectively. The vertical dashed line of 34.88 THz was closed to the maximum absorption finger located between 30 to 40 THz, at least within half-height width of the finger. **B, C:** The potential of mean force (PMF) of K^+^ and Na^+^ ions permeate through ion channels, before and during MIRS with frequency of 34.88 THz (blue and red lines), respectively. The field strength was E_0_=2.5 V/nm.

The protein structures of K^+^ and Na^+^ channels were tetramers, consisting of a single chain and include a narrow pore region (i.e. selective filter), playing a decisive role of the permeation efficiency of K^+^ and Na^+^ ions. Therefore, by considering the calculation requirements of quantum chemistry method, we further simplified the ion channel structure into a filter model. We identified the specific absorption modes of ion channels during MIRS (Supplementary Fig. S2) according to the filter structure extracted from the model of K^+^ and Na^+^ channels, the intrinsic spectrum was further calculated by using Gaussian 09 software based on density functional theory (DFT) at B3LYP/6-31G(d) level. It needed to be noted here that the absorption finger of − OH^−^ groups at the filter region of Na^+^ channels are distant from ∼34.88 THz (Supplementary Fig. S2), indicating that MIRS was mainly absorbed by − C=O groups at the inner wall of the filter region of K^+^ channels. These data demonstrated that the effect of MIRS with 34.88 THz could enhance the resonance absorption of K^+^ channels: the − C=O groups are frequency-sensitive due to their collective resonance to the K^+^ channels, while these effects seldom occurred in Na^+^ channels or water molecules.

In response to a signal, neurons emit action potentials mainly generated by the ion permeability events of sodium ions through Na^+^ channels and potassium ions through K^+^ channels. To compare the modulation of neuronal electrical activity by MIRS, we computed potential of mean force (PMF) as potential energy needed to cross configurations during ion permeability (Bernèche and Roux, 2001; Li, et al., 2021).By applying identical computational methods, the PMF before MIRS showed a higher potential energy at the entrance than at the exit of selectivity filter for both K^+^ channels and Na^+^ channels (blue lines in Fig. 5B and C). It implied that potassium ions and sodium ions could permeate the selectivity filter with comparable energetic potential. A further MD simulation showed that the potential energy difference of potassium ions through the filter became enlarged dramatically during MIRS, while the energy of sodium ions changed slightly through the filter configuration of Na^+^ channels (red lines in Fig. 5B, C). This indicated a selectively enhanced K^+^ permeability. To statically characterize the effect on ion permeability by MIRS, we defined the PMF ratio of ΔP_MIRS_/ΔP_before_ (ΔP_before_, ΔP_MIRS_ represent for the case before and during MIRS) as the change of ion permeability of potassium and sodium ions, respectively. Data showed that MIRS could increase the PMF ratio of K^+^ channels about 1.4-fold at the site around ξ = 1.1 nm, while the ratio of Na^+^ channels was kept closely to 1 at the site around ξ = 0.5 nm. Together, our simulation data demonstrated that MIRS with frequency of 34.88 THz could be selectively absorbed by K^+^ channels, and thus improved the efficiency of K^+^ permeability, while might not affect Na^+^ channels effectively. Therefore, MIRS could have a potential to alter neural discharge activities by selectively and significantly enhancing potassium permeability through K^+^ channels.

## Discussion

Our study provides several lines of evidence that MIRS could cause reversible and gain modulation on neuronal activity and sensorimotor behavior, by simultaneously recording neuronal and behavioral responses in awaking-behaving pigeons (Fig. 6). Our results showed that MIRS could achieve neuronal excitation and inhibition in the same pretectal neuron and cause gain modulation on pursuit eye movements of OKN. These alternations depended on the level of ongoing firing rates evoked by the directions and velocities of visual motion. Further computational simulations revealed that MIRS may effectively enhance K^+^ permeance through the selectivity filter of potassium channels (Fig. 5, 6B). Therefore, MIRS with 34.88 THz could be a specific neuromodulation approach to excite and inhibit neuronal firing and then alter sensorimotor behavior in a manner of the strength of sensory inputs.

**Figure 6.**
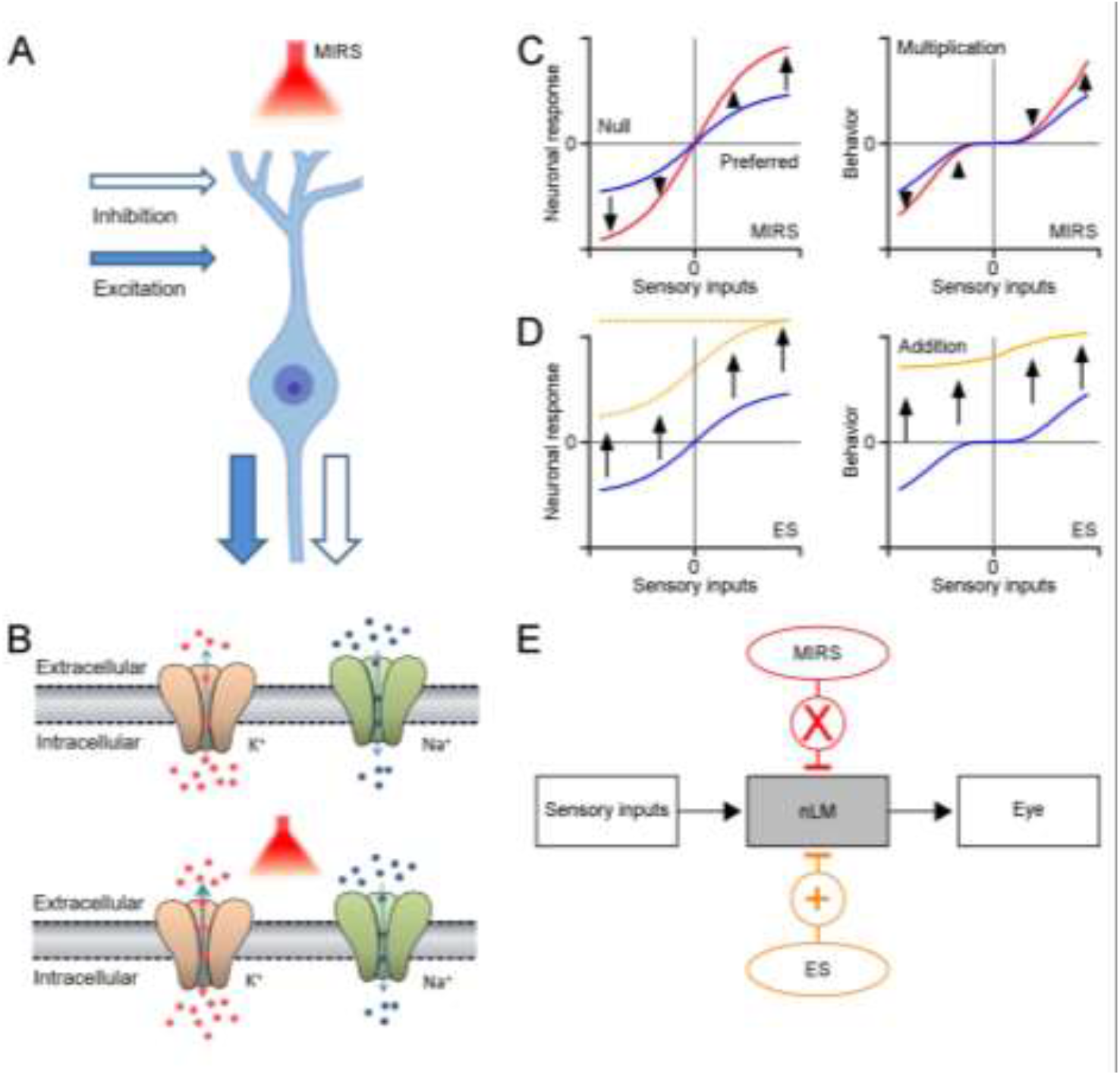
Schematic diagrams showing modulations of MIRS and ES suggested by our data. **A:** MIRS produces positive and negative modulations on visual responses in the same pretectal neuron. **B:** MIRS preferentially regulates permeation of K^+^ channels, instead of Na^+^ channels. **C:** MIRS exerts multiplicative gain modulations on neuronal responses to sensory and behavior performances, suggested by our experimental data. Blue and red lines present neuronal and behavior responses before and during MIRS. **D:** ES could exert additive modulations on neuronal firing, and cause unidirectional deflections in behaviour in our results. Blue and yellow lines present neuronal and behavior responses before and during ES. Dashed lines present the possible changes in neuronal firing by ES suggested by prior researches. **E:** Summary of different effects of MIRS and ES on sensorimotor transformation: gain modulation by MIRS, and additive modulation by ES.

OKN consists of pursuit eye movements during slow phases tracking in the direction of large-field motions and saccadic eye movements in fast phases resetting back into the opposite direction. OKN responses are highly sensitive to the velocity of visual motion. We found that MIRS applied in nLM induced gain modulations on pursuit eye movements based on the visual motion: MIRS can significantly speed up pursuit eye movements induced by a faster visual grating motion, but failed to significantly modulate pursuit when OKN was evoked by a slower visual grating motion. Note that if the grating was stationary (0 deg/s), pigeons only made spontaneous saccadic eye movements to search surrounding instead of pursuing visual motion, MIRS also failed to initiate any pursuit eye movements. Compared with MIRS, the classical ES in nLM can effectively deflect eye movements toward the T-N direction isolated from directions of grating motion. Even pigeons well pursued a grating motion in the N-T direction or freely searched still gratings without any pursuit, eye movements were deflected and driven to pursuit in the T-N direction immediately once ES applied. Comparison behavioral evidence from MIRS and ES, we found that MIRS could induce gain modulations to regulate oculomotor behavior depending on the strength of sensory inputs.

Neuronal circuitry for OKN has been studied for decades. In birds, the pretectal nLM plays complementary roles in encoding vision information of a large-field optic flow and guiding pursuit eye movements in slow phases of OKN (Cao, et al., 2004; Yang, et al., 2008b). Major neurons in the pretectal nLM encoded visual information of horizontal motion (Wylie and Frost, 1996; Wylie and Crowder, 2000). Visual responses of nLM were sensitively tuned by the velocity of grating motion. There were more than half pretectal neurons excited their visual responses to the T-N directional motion and inhibited their firing rates to the N-T directional motion. Once the grating moved from 2 deg/s to 8 deg/s in both preferred and null directions, firing rates of pretectal neurons could be activated at different visual responsiveness levels, emitting neuronal discharges into high- or low-frequency. These paved a fundamental basement to specifically investigate the neuromodulation effects of MIRS according to levels of neuronal visual responses. Our results showed that MIRS can further excite or inhibit neural firing occurred in the same neuron in a manner of its level of visual responses, respectively. Meanwhile, when animals viewed a given grating motion, the same MIRS could facilitate neuronal activity in one nLM neuron and suppress firing rates in the other neuron in a manner that the grating motion excited or inhibited neuronal discharges of that cell. Therefore, MIRS can cause reversible and multiplicative gain modulations on neuronal signals depending on the ongoing levels of visual responses or the strength of sensory inputs, which could follow the rule of “when strong is strong, when weak is weak” (Fig. 6C).

Traditional views thought infrared neuronal stimulation could cause local thermal heat by water absorption, which changes capacitance of the transmembrane to excite cells (Shapiro, et al., 2012), or activate thermosensitive TRPV channels to depolarize cells(Albert, et al., 2012). Recently, experimental evidence in *vitro* demonstrated that MIRS with a specific wavelength (5.6 μm) will cause nonthermal effect on ion channels and neuronal functions, especially when the distance was greater than 300 μm. Our evidence from behavioral and neurophysiological results are consistent with the nonthermal effect of MIRS. First, the distance between the MIRS fiber tip and recording sites was kept about 850 ± 300 µm (Fig.1B), which was limited thermal changes within 3°C(Tan, et al., 2021). Second, the modulation of pursuit eye movements by MIRS depends on the sensory inputs, which is not consistent with the thermal effect. MIRS increased pursuit velocities under a grating motion of 8 deg/s. If we assumed it was caused by local thermal heat of MIRS, then this heating effect should keep working to speed up oculomotor behavior when gratings move at lower velocities or keep still. Obviously, behavioral results showed a clear sensory-input depended modulation to against the thermal effect. Third, visual responses of the same recorded neuron can be increased by MIRS only when it was excited by motion in the preferred direction, and decreased by the same MIRS when it was inhibited by motion in the null direction. These positive and negative effects depended on the evoked excitative or inhibitive visual responses in individual neurons. While we are not able to eliminate the thermal effect induced neuronal activity changes or vasodilation as contributing factors, experimental evidence observed demonstrate consistently that the bidirectional and gain modulation of MIRS on neuronal responses and behavioral performances depends on the strength of sensory inputs (Fig. 6C).

The multiformity of ionic channels allows neurons to encode and transfer information by generating action potentials with a wide range of shapes, patterns and frequencies. This process must involve complex interactions with ion channels. A recent research in brain slices found that multiplicative modulations of the action potential generation by MIRS depended on the strength of current pules in *vitro*, and an increase in K^+^ currents (not Na^+^ currents) could lead to the gain modulation (Liu, et al., 2021). Consistent with previous reports, our ion channel simulations indicated that MIRS can cause the carbonyl group (-C=O) enriched on the K^+^ channel selective filter to resonate, thereby decreasing in input resistance and increasing in potassium ion flow, which could also work to modulate neuronal responses to sensory inputs in *vivo* (Fig. 6B). Potassium channels commonly playing a major part in the repolarization of action potentials. The increase in potassium ion flow by MIRS could cause a faster and/or earlier repolarization, and lead to a shortened action potential duration (Supplementary Fig. S3; (Liu, et al., 2021)) and an enlarged afterhyperpolarization. Once there were “stronger” stimuli (i.e. pretectal neurons were excited in the preferred direction in our case), these might cause faster recovery from inactivation of sodium channels (Bean, 2007) and the prior action potential’s refractory period. Then the neuron can be facilitated to initiate a subsequent action potential, which resulting in fasting of high-frequency firing. Once there were “weaker” stimuli (i.e. pretectal neurons were inhibited in the null direction), the raising K^+^ permeability might hinder the depolarization and retard the threshold potential of a subsequent action potential, which resulting in slowing of low-frequency firing. Although the computational simulations have not listed a full description from ion channels to generations of action potentials, our data provided evidence that suggest ionic mechanisms underlying MIRS: MIRS could preferentially modify permeation of K^+^ channels, leading to alternations of action potential generation in a manner of ongoing firing levels depended on sensory inputs to guide behavioural performances.

Distinct from the electrical stimulation that would activate neuronal firing and cause unidirectional deflections in behaviour (Fig. 6D), MIRS produces positive and negative modulations on visual responses in the same neuron, and then lead to gain modulations on behavioral performances (Fig. 6C-E). Distinct from the optogenetic stimulation that would require application of light-sensitive genes, our results showed that MIRS could selectively activate or inhibit neuronal responses by controlling the strength of sensory inputs in an individual or population cells, with gene free manipulation. Moreover, our results demonstrated that effects of MIRS occur reversibly in neuronal and behavioral responses in awake-behaving animals. These findings together suggested that MIRS could be used as a promising neuromodulation approach to excite and inhibit neuronal firing in brain researches and clinical applications.

## Methods

### Animal preparation

We conducted experiments on 16 awake, behaving adult pigeons of either sex (*Columba livia*, body weight: 300-500g). 10 of animals contributed for MIRS experiments combining recording in the pretectal nLM, while 6 of them were used for electrical stimulation experiments. We performed experiments using techniques that have been described in detail before(Yang, et al., 2008a; 2008b; Yang, et al., 2017). Procedures were in accordance with the guidelines for the care and use of animals established by the Society for Neuroscience and approved by the *Institutional Animal Administration Committee* at the Institute of Biophysics, Chinese Academy of Sciences.

### Visual conditions

A large-field square wave grating was generated by MATLAB program and projected to a screen subtended a visual field of 130° by 140°. The grating consisted equal-width black and white stripes with spatial frequency of 0.16 cycles/degree. The meridians of the visual field were rotated by 38° (Britto, et al., 1990; Fu, et al., 1998) to match pigeons’ normal viewing conditions (Erichsen, et al., 1989). Animals produced saccades and OKN when they viewed stationary gratings (0 deg/s) and moving gratings. The grating moved at 2 deg/s and 8 deg/s in the T-N and N-T directions to elicit OKN, respectively (Gioanni, et al., 1983; Yang, et al., 2008b).

### Mid-infrared stimulation and Electrical stimulation

The MIRS was performed using a quantum cascade mid-infrared laser (Daylight solution Inc., model MIRcat) with a radiation wavelength of 8.6 μm, frequency of 34.88 THz, and a power of 80 mW. The pulse train was applied for 120 seconds with a mark of 2 μs and a space of 3 μs. An infrared fiber (IRF-S-9, IRflex) with a diameter of 600 μm was used for coupling. The fiber was inserted vertically into the left pretectal nLM. The distance between recording microelectrodes and fiber tips was about 850 ± 300 µm (mean ± SD, Fig.1 B). There were ∼10 minute intervals between any two MIRS to get a recovery from the prior stimulation.

The ES was generated by an isolated pulse stimulator (A-M Systems, model 2100) and applied for 120 seconds in the pretectal nLM, with parameters of 0.2 mA, 100 μs pulse width, 133 Hz(Elias, et al., 2020). The stimulation electrodes were glass-insulated tungsten bipolar electrodes with an exposed tip of 60 µm and a distance of 400 µm between the two tips. The electrodes were advanced into the brain following the same method as optical fibers in the MIRS.

### Data acquisition and analysis

#### Extracellular recording

We introduced homemade glass-insulated tungsten microelectrodes into the pretectal nLM, with an impedance of 1-3 MΩ (Gioanni, et al., 1983; Yang, et al., 2008b). We amplified extracellular action potentials, filtered them with a bandpass of 300 Hz to 5 kHz (A-M Systems, Model 1800), and digitized at 25 kHz (CED, Power 1401 Cambridge electronic design limited,) for off-line spike sorting (Spike2, Cambridge electronic design limited). At the end of each experiment, recording sites were marked by electrical destruction (positive current of 30-40 µA for 20-30 seconds).

#### Electrooculogram recording (EOG)

Eye position changes were recorded by an electrooculogram system (Wohlschläger, et al., 1993; Yang, et al., 2008a; 2008b), sampled at 2.5kHz and stored simultaneously with neuronal signals. EOG signals were smoothed by a low-pass filter with a cut-off frequency of 5 Hz, and differentiated into eye velocity. For calibration purposes, eye movements were video graphed by an infrared video camera simultaneously with EOG recording (Niu, et al., 2006; Yang, et al., 2008a; 2008b). Eye movement data were defined as positive values for T-N direction and negative values for N-T direction.

Eye movements and neuronal spikes were calculated in an interval from 0-600 ms after the onset of pursuit eye movement in OKN. The choice of the interval ensures that our measures are related to the visual response to grating motions during pursuit of OKN. The onset and offset of pursuit eye movement were determined with a customized MATLAB code according to characteristic oscillations in avian saccadic eye movements, and then manually rechecked. Data were excluded if the period of pursuit was shorter than 600 ms, although this probability was negligible under tested visual conditions (Fig. 1E). Neuronal activities were collected before, during and after MIRS for about 120 seconds, and smoothed by a Gaussian Kernel filter with an *h* value of 25. We defined neuronal firing to be facilitated or suppressed by MIRS if they were significantly higher or lower than activities before MIRS (Wilcoxon signed-rank test, p<0.05).

### Molecular dynamics simulation

The simulation was carried out to obtain an understanding of the power of ions permeation, and aimed to compare the permeability of K^+^ and Na^+^ ions during MIRS at nano-scale spatial and femto-second time resolution. Based on the eukaryotic model of voltage-gated K^+^ channels (PDB ID: 2a79, Supplementary Fig. S1) (Long, et al., 2005) and Na^+^ channels (PDB ID: 4dxw, Supplementary Fig. S2) (Zhang, et al., 2012), respectively. In our simulations, the force field of CHARMM 36 and periodic boundary conditions were used (Mackerell and Nilsson, 2008). The connection element algorithm Ewald was used to deal with the electrostatic interaction (Leeuw, et al., 1980). The Velocity-Verlet algorithm (Andersen, 1983) was performed to solve the motion equation, with the time step of 2 fs. All bond lengths were limited by the Lincs algorithm (Swope and William, 1982). In particular, for K^+^ channels, the truncation of Lennard-Jones interaction and the real space part of Ewald sum were 1.90 nm, the convergence factor of Ewald sum was 1.65 nm, and the radius of K-space section was 10.4 nm. Meanwhile, for Na^+^ channels, the truncation of Lennard-Jones interaction and the real space part of Ewald sum were 1.623 nm, Ewald and convergence factors were 1.65 nm, and the radius of K-space section was 10.4 nm. The process of the ion permeation events was divided into two stages according to before and during MIRS. First, we fixed the phospholipid bilayer and protein (except the filter region of K^+^ and Na^+^ channels) in the system to research the process of ion permeation through the filter domain under the NVT ensemble at room temperatures 300 K. Second, to combine effects of the electromagnetic wave on the ion channel, MIRS with frequency of 34.88 THz was added into the whole simulation system (Wu, et al., 2020). The intensity ratio of the electromagnetic component of an electromagnetic wave is equal to the speed of light. In the formula:

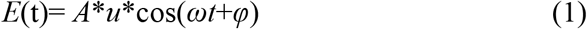

where *A* represented the maximum amplitude of the electric field to determine the electric field strength of electromagnetic wave, and *u* and *φ* represented the polarization direction and phase, thus set to (0, 0, 1) and 0 respectively. The electromagnetic wave frequency was computed as a function of the angular frequency ω by the equation *γ* = *ω*/2π. The total time was ∼10 ns, and the trajectory is saved every 0.1 ps for subsequent data analysis.

We simulated the absorption spectrum of K^+^ and Na^+^ channels by using molecular dynamics methods based on the classical GROMACS package (Abraham, et al., 2015)(Fig. 5A). The absorption spectrum was calculated according to the Fourier transform of the velocity autocorrelation function of the total charge current of our simulation systems (Heyden, et al., 2010). We set the time interval of spectrum sampling as 1 fs, and the total time of sampling as 50 ps. The absorption spectra were calculated based on the Fourier transform of the autocorrelation function of the total charge current (Heyden, et al., 2010):

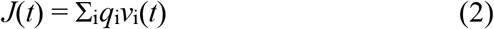

where *q*_i_ represented the charge of the i-th atom, and *v*_i_(*t*) stood for the velocity of the i-th atom at time *t*.

We further simulated the potential of mean force (PMF) as potential energy needed to cross configurations during ion permeability (Bernèche and Roux, 2001; Li, et al., 2021)(Fig. 5B and C). Umbrella sampling (US) is a method that a series of initial configurations are sampled along a reaction coordinate defined between two groups, then we simulated in the group of K+/Na+ harmonically restrained against the other fixed group via an umbrella biasing potential. Initially, a K^+^ or Na^+^ ion was placed in the z direction at the entrance of the selectivity filter. The Cl^-^ ion was in line with the center of the protein channel. It was initially frozen and taken as the reference group. The simulated K^+^ or Na^+^ ion was pulled by a force constant of 2000 kJ/mol/nm^2^ along the filter for 1.5 ns at a pulling rate of 0.01 Å per ps. All residues of filter were set flexible during pulling. The initial configurations for the US simulations were extracted from the pulling process at an interval of approximately 0.2 Å (COM distance between the reference Cl^-^ ion and the simulated K^+^ or Na^+^ ion) along the ion conductance path. Pressure equilibration was carried out for 1.0 ns for all sampled windows, followed by 3.0 ns trajectory generation for a reliable PMF calculation. In each umbrella sampling simulation, the simulated ion was restrained harmonically with a force constant of 5000 kJ/mol/nm^2^ along the z-axis. During the US process, the protein (except for the key SF residues) was restrained, so the shift of the whole protein due to system thermal and pressure fluctuations can be negligible. The free energy profile was calculated with the WHAM method implemented in GROMACS based on the US data.

In general, the critical factor to determine the permeability of ions concern to the filter region of ion channels (Kopec, et al., 2018). The selective filter of K^+^ channels composed of 24 residues (sequence index of 75∼80) (Long, et al., 2005), and 12 residues for Na^+^ channels (sequence index 9∼11) (Zhang, et al., 2012). According to the filter structure extracted from the model of K^+^ and Na^+^ channels, the intrinsic spectrum was further calculated by using Gaussian 09 software (Frisch, et al., 2009) based on density functional theory (DFT) at B3LYP/6-31G(d) level (Zhang, et al., 2012), thus to explore the specific absorption modes of ion channel during MIRS (Supplementary Fig. S2).

## Acknowledgments

We thank Xi Xu and other members of our laboratory for helpful comments on an earlier version of the manuscript and discussions. We thank Qian Wang, Chen Wu, and Haiyan Liu provided invaluable technical assistance.

## Supplementary Figures

**Figure S1.**
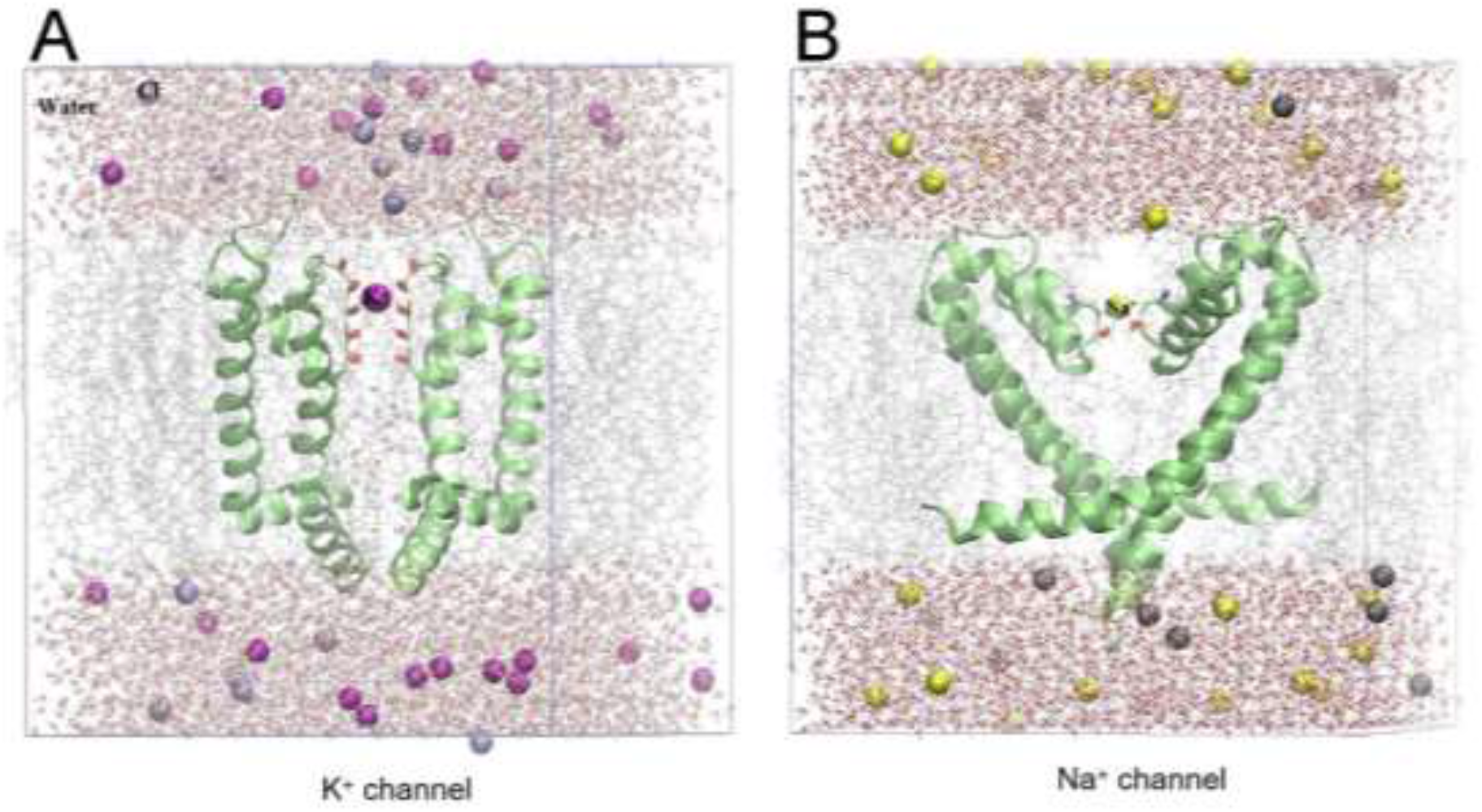
The composite atomic models contain the K^+^ channel (A) and the Na^+^ channel (B). In our models, we constructed channels containing the whole protein placed at the middle of phospholipid bilayer to separate water and ions on each side. There are 12 potassium, 12 sodium ions and 20 chloride ions for K^+^ channels, and 14 sodium ions, 14 potassium ions and 24 chloride ions for Na^+^ channels, respectively. Both the water thickness of K^+^ and Na^+^ channels are ∼0.3 nm. **A:** The K^+^ channel (PDB ID: 2a79) contains 10,294 atoms, including 2,725 TIP3P water molecules, 12 K^+^ and 20 Cl^-^ ions, thus the concentration of a salt solution is 0.15 M. The protein has four negative charges, the addition of four counterbalance ions ensures that the whole system is electrically neutral. The size of the PBC box is 5.04 nm × 5.16 nm × 6.25 nm. **B:** The Na^+^ channel (PDB ID: 4dxw) contains 14,922 atoms, including 4,067 water molecules, 14 Na^+^ and 24 Cl^-^ ions, thus the concentration of a salt solution is 0.15 M. Similarly, the addition of four counterbalance ions also ensures that the whole system is electrically neutral. The size of the PBC box is 5.40 nm × 5.40 nm × 7.30 nm. The purple atom represents K^+^, the yellow atom represents Na^+^, and the gray atom represents Cl^-^. The TIP3P water molecule is displayed in the format of a red ball stick. Green strips show K^+^ and Na^+^ channels, red stripes show − C=O groups located at the inner wall of the filter region, blue stripes show -OH^-^ groups of the filter, and light gray lines show a phospholipid bilayer, respectively.

**Figure S2.**
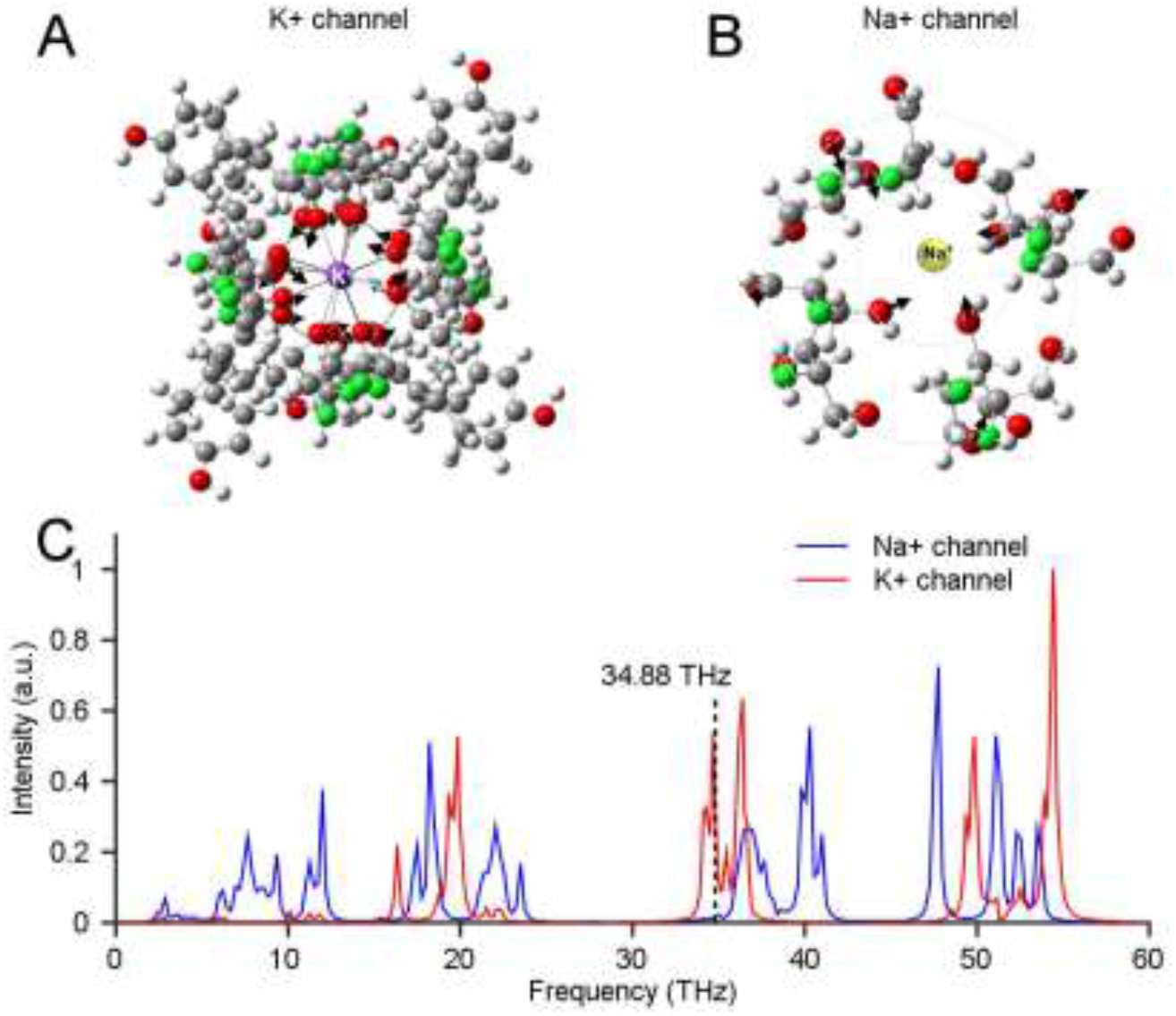
The eigen-modes and intrinsic spectrum calculation for K^+^ and Na^+^ channels. **A and B:** Calculations on the eigen-mode frequencies of –C=O groups for K^+^ channels, and –C=O, –OH^−^ groups for Na^+^ channels according to the filter structure extracted from the model of K^+^ channels (PDB ID: 2a79) contain 284 atoms (**A**) and Na^+^ channels (PDB ID: 4dxw) contain 96 atoms (**B**). **C:** The intrinsic spectrum was calculated by using Gaussian 09 software based on density functional theory (DFT) methods at B3LYP/6-31G(d) level.

**Figure S3.**
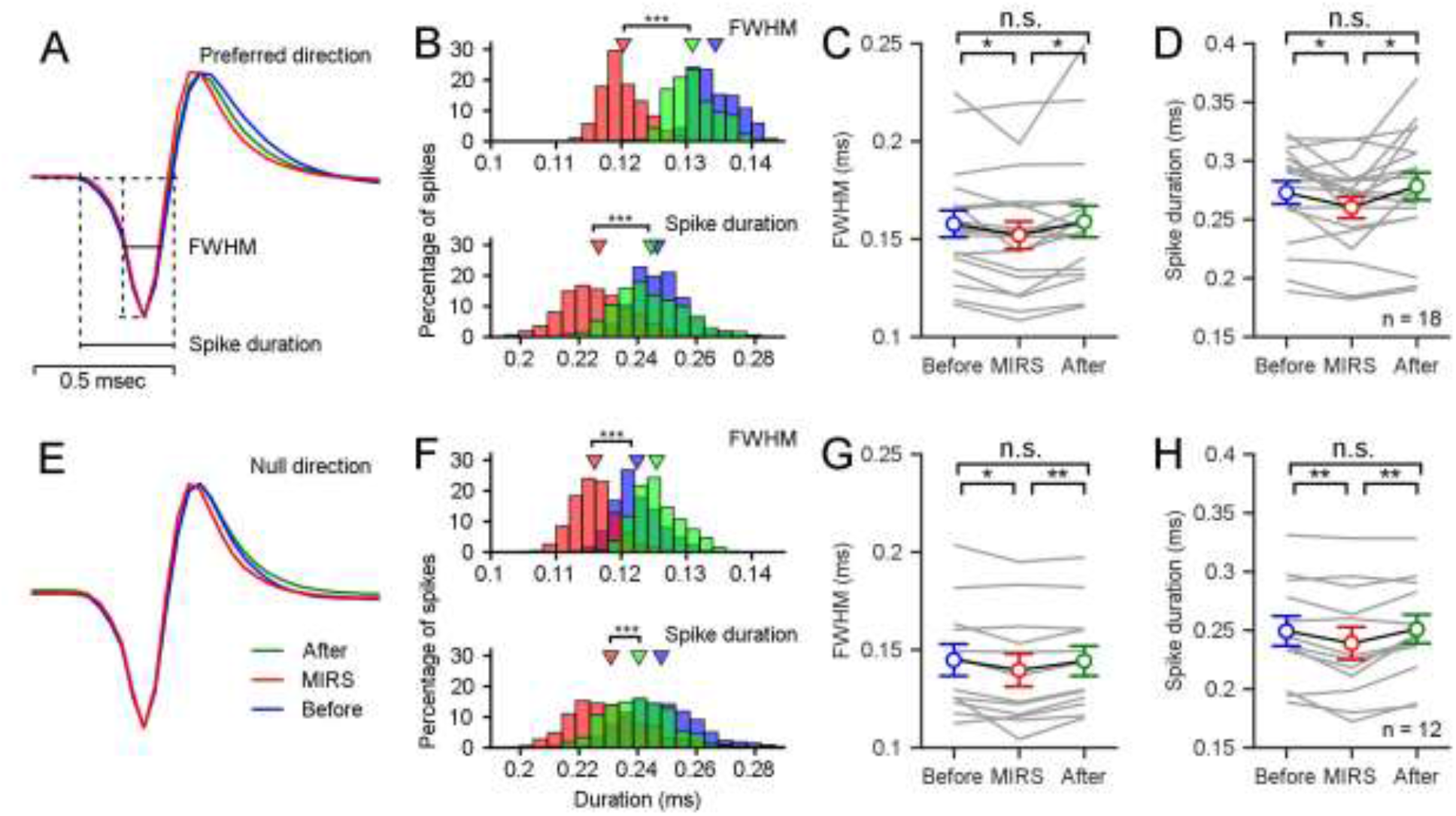
Extracellar recordings showed shortened action potential durations of pretectal nLM neurons during MIRS in the preferred direction (A-D) and the null direction (E-H). **A** and **E**: mean action potential traces from an example pretectal neuron before, during and after MIRS in both directions. **B** and **F**: Distributions of the full width at half maximum (FWHM) and the duration of spikes’ negative phases (Spike duration) of the example pretectal neuron’s action potentials (Student’s T-test, *** P<0.001). **C, D, G**, and **H**: Statical summarization of the full width at half maximum and the duration of spikes’ negative phases across population neurons (n=18 in **C** and **D**; n=12 in **G** and **H**; Paired Student’s T-test, * P<0.05, ** P<0.01; Error bars represent 1 SEM). In order to reduce the influence of recording noises on the definition of durations, we analyzed data from recorded neurons with the signal-to-noise ratio above 4:1 in Figure 4A and B.

